# A multifunctional *Dehalobacter*? Tandem chloroform and dichloromethane degradation in a mixed microbial culture

**DOI:** 10.1101/2023.08.10.552028

**Authors:** Olivia Bulka, Jennifer Webb, Sandra Dworatzek, Radhakrishnan Mahadevan, Elizabeth A. Edwards

## Abstract

Chloroform (CF) and dichloromethane (DCM) contaminate groundwater sites around the world, which can be remediated through bioaugmentation. Although several strains of *Dehalobacter restrictus* can reduce CF to DCM, and multiple Peptococcaceae can ferment DCM, these processes cannot happen simultaneously due to CF sensitivity in the known DCM-degraders or electron donor competition. Here we present a mixed microbial culture that can simultaneously metabolize CF and DCM to carbon dioxide and create an additional enrichment culture fed only DCM. Through species-specific qPCR, we find that a *Dehalobacter* strain grows both while CF alone and DCM alone are converted, indicating its involvement in both metabolic steps. Additionally, the culture was maintained for over 1400 days without addition of exogenous electron donor, and through electron balance calculations we show that DCM mineralization produces sufficient reducing equivalents (likely hydrogen) for CF respiration. Together, these results suggest intraspecies electron transfer could occur to continually reduce CF in the culture. Minimizing the addition of electron donor reduces the cost of bioremediation, and understanding this mechanism informs strategies for culture maintenance and scale-up, and benefits contaminated sites where the culture is employed for remediation worldwide.

**SYNOPSIS:** Dechlorination of chloroform to dichloromethane and dichloromethane mineralization are performed concurrently by a *Dehalobacter*-containing mixed microbial community without provision of exogenous electron donor.

**TOC ART:** 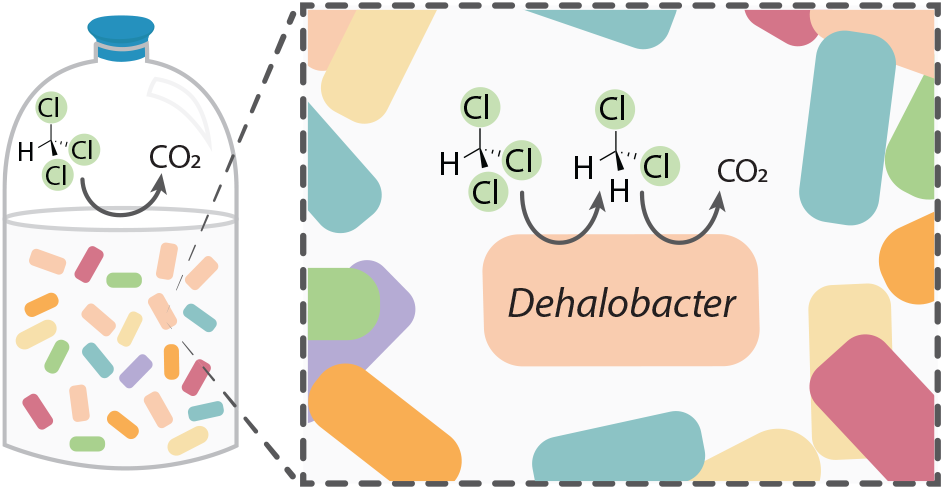

## INTRODUCTION

Chloroform (CF) and dichloromethane (DCM) are widely used halogenated solvents that have significant environmental impacts due to their toxicity, persistence, and potential for bioaccumulation. These pollutants contaminate soil and water resources—more than 717 CF-contaminated sites are on the EPA’s National Priority List in the United States alone—and pose a threat to human health and ecosystems (1). Bioremediation through bioaugmentation has emerged as a promising strategy for the removal of these compounds from the environment.

Several *Dehalobacter restrictus* strains respire CF to DCM through reductive dechlorination (2–6). These strains derive energy using CF as an electron acceptor, and typically require hydrogen or formate as an electron donor. This need for reducing equivalents necessitates injecting sometimes expensive, electron-rich materials to stimulate dechlorination *in situ* (7). DCM degradation is a critical second step to CF bioremediation in contaminated groundwater sites, but in many CF-dechlorinating cultures the dechlorination process often halts with DCM accumulating in the system, preventing further CF biodegradation (2, 4). Furthermore, though some DCM-degrading microbes have been identified, most are very sensitive to CF, rendering their use for bioremediation in complex contaminated sites impractical (8–13).

Two DCM degradation pathways have been reported in literature (Figure 1): 1) DCM fermentation—as performed by *Dehalobacterium formicoaceticum* and *Candidatus* Formimonas warbariya (8, 10, 14, 15)—and 2) DCM mineralization—as performed by *Candidatus* Dichloromethanomonas elyunquensis (9, 16). The former, DCM fermentation, consumes carbon dioxide and uses the Wood-Ljungdahl pathway to produce acetate and formate through internally balanced redox reactions (8, 14). DCM mineralization also likely uses the Wood-Ljungdahl pathway, but produces carbon dioxide and hydrogen as oxidized and reduced products (17). This requires syntrophic microbial community members, including methanogens and acetogens, to consume the accumulating hydrogen and maintain sufficiently low concentrations for additional mineralization to occur (17).

**Figure 1.**
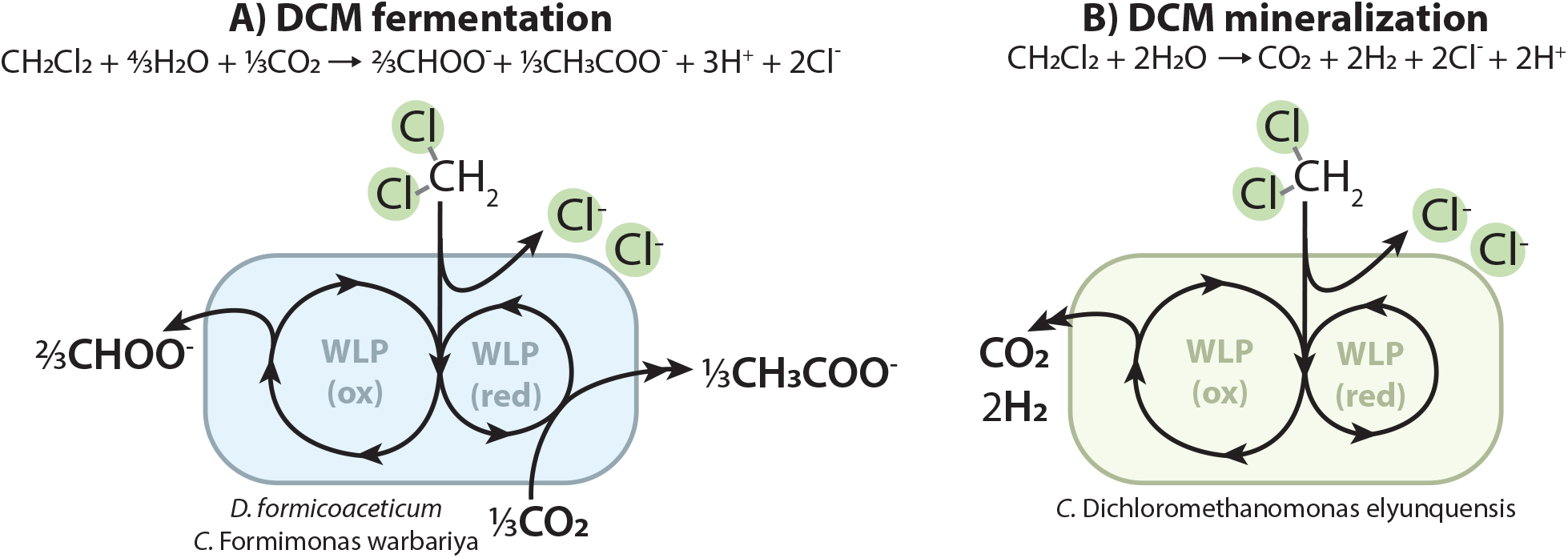
Schematic of the two known DCM-consuming metabolic pathways: **A)** DCM fermentation compared to **B)** DCM mineralization, and the microorganisms known to perform each process. Adapted from Chen *et al*. (17).

SC05, sold commercially as KB-1® Plus CF, is a stable anaerobic microbial enrichment culture originally derived in 2010 from a site in California contaminated with both trichloroethene (TCE) and CF (18). This culture can degrade both CF and DCM and has long been thought to contain two key organisms, one performing CF dechlorination and the other DCM fermentation in tandem. SC05 rarely accumulates DCM and a previous study of the culture used ^14^C-labelled CF to demonstrate its transformation to DCM, followed by conversion to carbon dioxide (18). The oxidation of DCM to CO_2_ is consistent with the DCM mineralization pathway—as studied in *C*. Dichloromethanomonas elyunquensis—which also results in production of hydrogen (17). This early work suggests the possibility of SC05 to produce its own electron donor from the mineralization of DCM to reducing equivalents usable by the CF-dechlorinator, but there have been no in-depth studies of the culture’s microbial composition or DCM-degrading capabilities published to date.

Here we **1)** describe circular electron shuttling (or self-feeding) in a derivative of the SC05 culture stimulated by the mineralization of DCM, and **2)** implicate *Dehalobacter restrictus* in both CF dechlorination and DCM mineralization—the first *Dehalobacter* seen to diverge metabolically from organohalide respiration.

## MATERIALS & METHODS

### Microbial cultures and growth conditions

The original SC05 culture (also known as KB-1® Plus CF) has been maintained since 2010 at SiREM (Guelph, ON) in defined FeS-reduced, bicarbonate-buffered mineral medium (19) amended weekly with CF as the electron acceptor and methanol (1 mM) and ethanol (0.5 mM) as electron donors, each at an electron equivalency of 6× CF demand (2 electron equivalents (eeq) per mole of CF). A culture aliquot was provided to the University of Toronto in 2018, referred to herein as SC05-UT, where it was initially amended with CF, ethanol, and lactate (20). Subsequently, the culture was only refed CF—without electron donor—due the work by Wang *et al*. showing that hydrogen production from DCM metabolism might be produced to fuel dechlorination without exogenous donor addition (18).

After maintenance for more than one year without electron donor amendment, a subsample of SC05-UT was inoculated into fresh medium (Day 0, Figure 2), which was named “DCME” (for DCM Enrichment) and henceforth maintained by addition of DCM alone. This culture was transferred twice more into 1 L of fresh medium at a 1% and 15% dilution, respectively, for further enrichment (Figure 2). At three points on the enrichment timeline (pink squares, Figure 2), sub-transfers of DCME and SC05-UT were used for specific time course experiments, as described below. Before each of these time course experiments, the culture was purged of volatiles with N_2_/CO_2_ (80:20, v/v) gas mix before inoculation to fresh medium in sterile blue butyl rubber stopper-sealed 250 mL glass serum bottles. The inoculations were performed in an anaerobic chamber (Coy) supplied with a N_2_/H_2_/CO_2_ gas mix (80:10:10 v/v).

**Figure 2.**
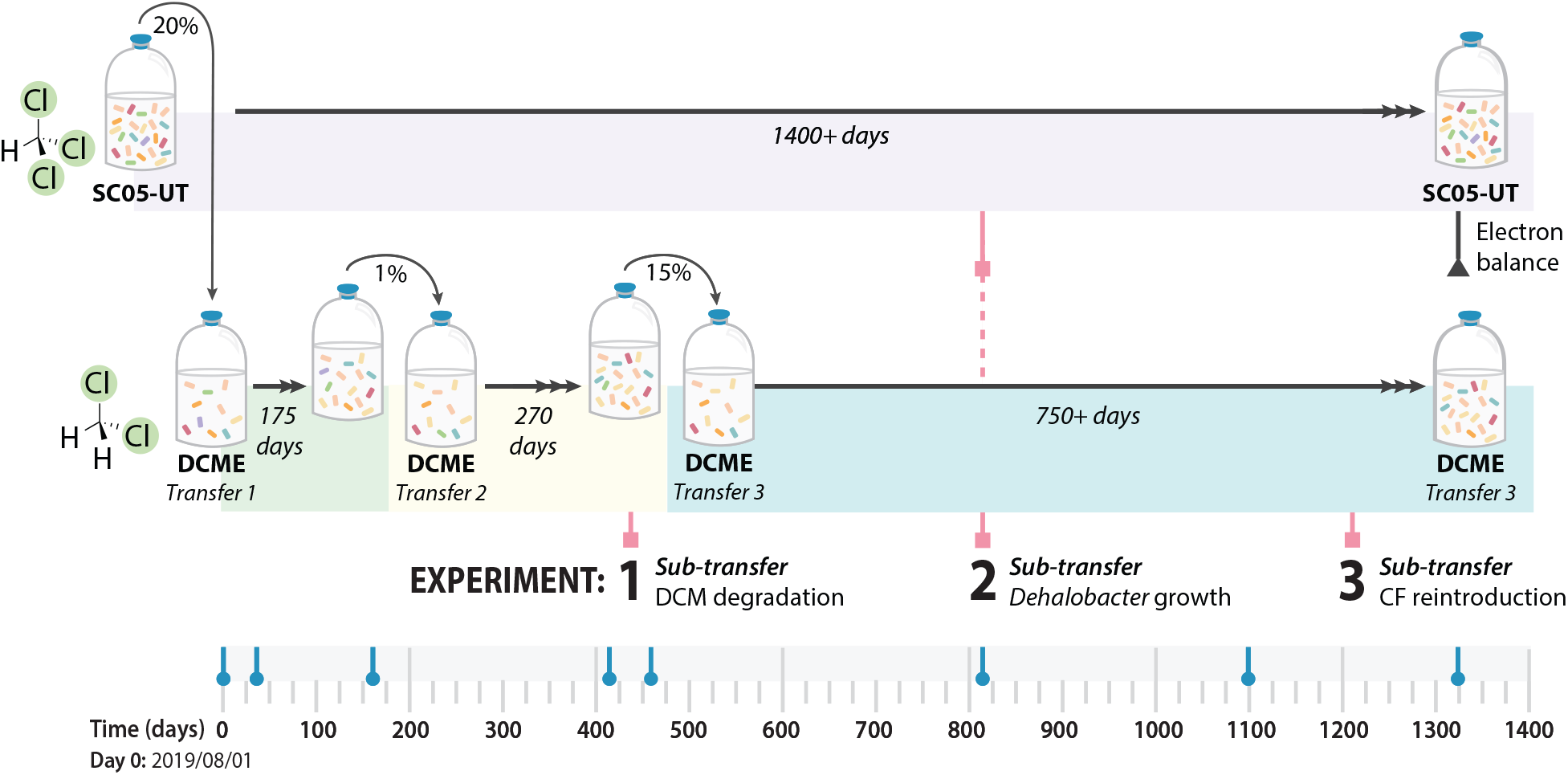
Timeline of SC05-UT and derived DCME transfers showing incubation periods over 1400 days. Sub-transfer dates for time course experiments #1-3 are marked with pink squares, and the date of electron balance measurements in SC05-UT is marked with a grey triangle. DNA sampling dates for both cultures are marked with blue circles.

### Time course experiment #1: DCM degradation study (Figure S3)

To determine the end products of DCM degradation and culture community composition of DCME, Experiment #1 was conducted after 433 days of enrichment on DCM. Three 25 mL aliquots were sub-transferred from the DCM enrichment culture to 150 mL fresh medium in 250 mL serum bottles and amended with 150 μmol DCM (0.85 mM aqueous) using a glass syringe. Three negative controls were prepared using autoclaved SC05-UT as the inoculum and spiked with 150 μmol DCM. Every 3-5 days, the headspace of each sub-transfer was sampled for gas chromatography analysis, and 1 mL aqueous samples were taken for organic acid ion chromatography analysis and DNA extraction followed by amplicon sequencing.

### Time course experiment #2: Dehalobacter growth study (Figure 5)

To establish the relationship between *Dehalobacter* and both CF and DCM transformation, *Dehalobacter* growth was quantified over one feeding cycle in subsamples of SC05-UT and DCME after 817 days of enrichment on DCM. Sub-transfers of SC05-UT (n=3) and DCME (n=2) were created by adding 15 mL of culture to 285 mL of fresh medium in 600 mL serum bottles and were fed 200 μmol CF (0.6 mM aqueous) and DCM (0.55 mM aqueous), respectively, with a glass syringe. Two negative controls were prepared using autoclaved SC05-UT as the inoculum and spiked with 150 μmol CF and 200 μmol DCM. The headspace of each sub-transfer was sampled for gas chromatography analysis every 3-5 days, and 1 mL aqueous samples were taken periodically for subsequent DNA extraction and qPCR. Each DNA sample was also sent for 16S amplicon sequencing to determine culture composition, which was scaled to the qPCR data. Aqueous samples were also taken from the SC05-UT transfers for organic acid analysis through ion chromatography.

### Time course experiment #3: CF dechlorination in DCME (Figure 6)

To determine whether DCME loses the ability to dechlorinate CF after >1200 days without exposure to CF, two additional sub-transfers were created by adding 5 mL of DCME to 95 mL medium in 160 mL serum bottles and amended with 25 μmol CF (0.25 mM CF aqueous) and 10 μmol hydrogen (1 mL H_2_/CO_2_, 20:80 v/v) to jump-start degradation. Two negative controls were prepared using autoclaved DCME as the inoculum and spiked with 25 μmol CF. The headspace of each bottle was sampled for gas chromatography analysis every 3-5 days, and 1 mL aqueous samples were taken for DNA extraction and qPCR weekly.

### Analytical procedures

CF, DCM, and methane were quantified by injecting a 0.3 mL headspace sample into a Hewlett-Packard 5890 Series II gas chromatograph (GC) fitted with a GSQ column (30-m-by-0.53-mm [inner diameter] PLOT column; J&W Scientific, Folsom, CA). The GC carrier gas pressure was set initially to 100 kPa, and the oven temperature was programmed to hold at 50°C for 1.5 min, then increase to 180°C at 60°C/min and hold for 2.5 min. Calibration was performed using external standards.

For organic acid analysis, 500 μL of culture was removed from a serum bottle using a 1 mL syringe in an anaerobic chamber (Coy) and filtered through a 0.1 μm pore syringe filter (Millipore). The flow-through was collected in a plastic microcentrifuge tube and stored at -80°C before analysis. The concentrations of lactate, acetate, propionate, pyruvate, oxaloacetate, malate, succinate, alpha-ketoglutarate, fumarate, citrate, and isocitrate were determined using a Dionex ICS-2100 ion chromatography system (Thermo Scientific) equipped with a Dionex™ IonPac™ AS11 IC column (Thermo Scientific) and a conductivity detector. Each sample was diluted 1/10 to a final volume of 500 μL, and 20 μL was injected onto the column incubated at 30°C at a flow rate of 1 mL/min, using a multistep KOH eluent gradient. The initial eluent concentration was 0.5 mM, increasing to 2.5 mM in 10 min, then increasing to 30 mM in 19 min, holding for 11 min, and then decreasing back to 0.5 mM in 0.5 min. Calibration was performed using external standards.

### DNA extraction and amplicon sequencing

DNA was extracted from 1 mL samples by pelleting the culture via centrifugation at 13000 x g for 15 minutes, then using the KingFisher™ Duo Prime (Thermo Scientific, Waltham, MA) and MagMAX Microbiome Ultra Nucleic Acid Isolation Kit to extract from the pellets as specified in the manual (Applied Biosystems, Waltham, MA).

DNA samples were sent for amplicon sequencing of the V6-V8 16S rRNA region using staggered primers (Table S1) by the Genome Quebec Innovation Centre using an Illumina MiSeq platform (Illumina Inc., San Diego, CA) and analyzed using an established QIIME 2 pipeline (21, 22). Briefly, the sequencing primers were trimmed from the data, then Dada2 was used to denoise and dereplicate sequences, remove chimeras, merge reads, and cluster them into features. The forward and reverse reads were truncated to 260 bp and 240 bp, respectively, and the maximum error rate was set to 2. Amplicon sequence variants (ASVs) were classified using the Silva database (v138). Archaea and bacteria communities are visualized separately to account for domain-specific biases.

### Quantitative PCR (qPCR)

Real-time quantitative PCR (qPCR) assays were performed to track *Dehalobacter* proliferation using 16S rRNA gene primers specific to *Dehalobacter restrictus* (F: 5’-AGCATTGGGTGTTAGGCGAA-3’; R: 5’-GCGCTAGCATTTTCAGAGGC-3’). The qPCR reaction mixtures were prepared in a UV-treated PCR cabinet (ESCO Technologies, Hatboro, PA), and contained 10 μL of 2× SsoFast EvaGreen® (Bio-Rad, Hercules, CA), forward and reverse primers (0.25 μM each, Table S1), and 2 μL of template DNA. The amplification program and analyses were conducted using a CFX96 Touch Real-Time PCR Detection System and the CFX Manager software (Bio-Rad). The qPCR method included an initial denaturation step at 98°C for 2 min, followed by 40 cycles of 5 s at 98°C and 10 s at 60°C, including a 2°C/s ramp between temperatures. Quantification was performed using 10-fold serial dilutions of PCR-produced standard DNA amplified from a larger region of the *Dehalobacter* 16S rRNA gene (F: 5’-CTGATCGTCGCCTTGGTAGG-3’, R: 5’-AGCAAGCAACATAAGCGAGTG-3’). The number of 16S rRNA copies/ mL of culture was calculated assuming a 100% DNA extraction efficiency.

### Electron balance calculations

Electron balances were calculated using measured concentrations of CF, DCM, methane (GC), acetate (IC), and *Dehalobacter* yields (qPCR) in SC05-UT and DCME. For each feeding cycle, final concentrations of CF, DCM, methane, and acetate were compared to initial concentrations to determine the amount of each compound consumed or accumulated in mmol, and converted to meeq by using the following conversion ratios: CF: 2 eeq/mol, DCM: 4 eeq/mol, methane: 8 eeq/mol, acetate: 8 eeq/mol. DCM mineralization was calculated by subtracting the amount of DCM accumulated from the amount of DCM theoretically produced from CF dechlorination. The fraction of donor electrons transferred to biomass was calculated using qPCR yields (Table S2) as previously described, using yields calculated in Table S3 (23).

## RESULTS & DISCUSSION

### SC05-UT dechlorinates CF and DCM without addition of an external electron donor

Though previous work suggested the possibility for CF dechlorination by SC05 without added electron donor (18), this is the first long-term report of this behavior without addition of methanol, ethanol, lactate, or exogenous DCM. Here, we maintain SC05-UT for over 800 days by re-amending CF alone over the entire period (Figure 3). Despite somewhat irregular refeeding due to limited lab access during COVID, in addition to other time constraints such as holidays, the culture retains consistent activity. Though we see some DCM buildup in the bottle from time to time, it is never stoichiometric to CF, suggesting concurrent degradation of both substrates—either by a single organism, or by two microorganisms working closely together by transferring electron-rich metabolites between them. No chloromethane was detected in the culture’s headspace, and no quantifiable organic acid accumulation was detected in this work or past work (18).

**Figure 3.**
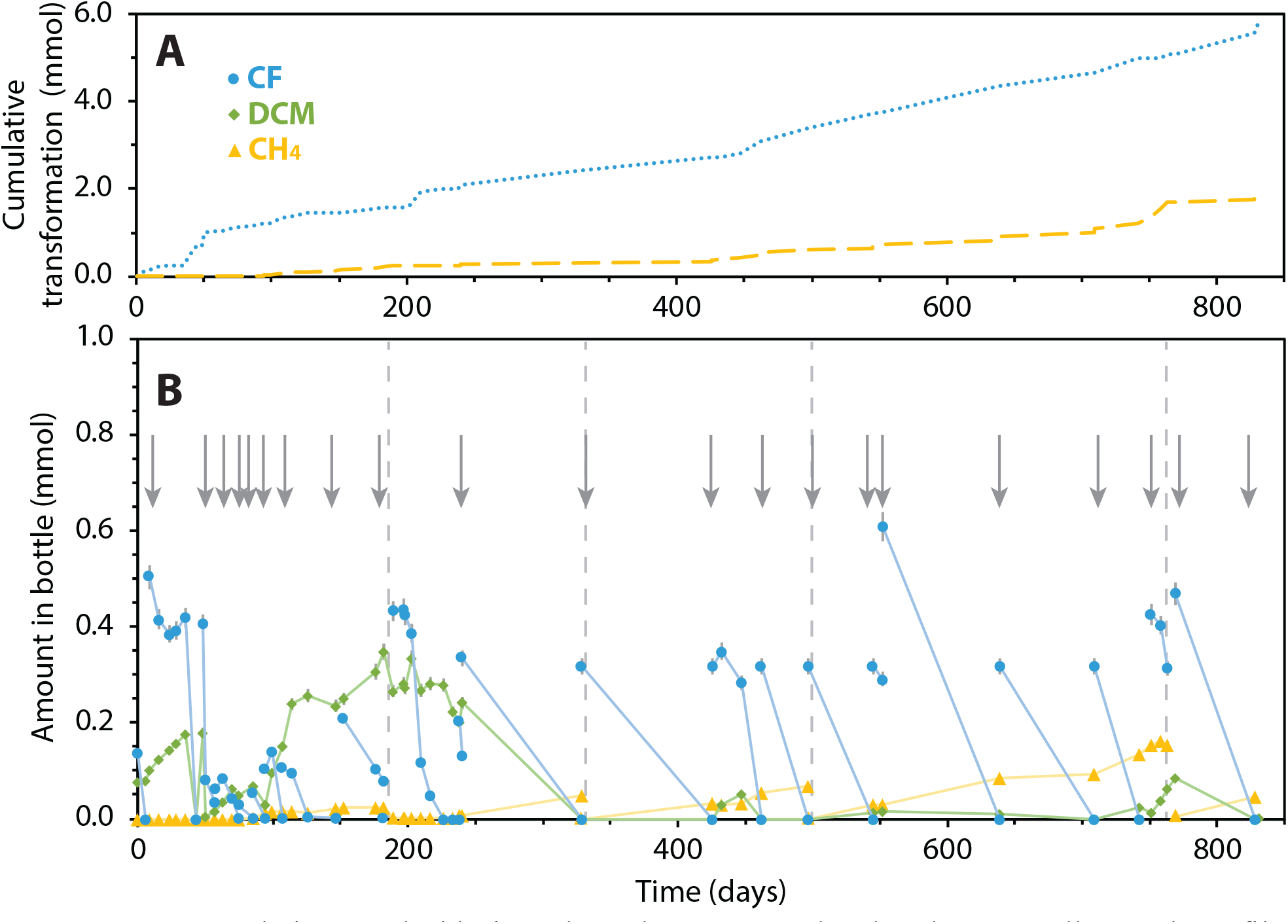
**A)** Cumulative CF dechlorinated, methane accumulated and **B)** overall growth profile by SC05-UT over 828 days (Days -15–813, Figure 2). Arrows indicate refeeding of CF, and grey dashed lines denote headspace purging of methane from bottle (n=1, error bars indicate analytical uncertainty of each measurement).

### Electron balance consistent with DCM mineralization in SC05-UT

To investigate the electron balance of SC05-UT, CF, DCM, and methane were tracked more closely over 3 CF feedings (Day 1327, Figure 2). As CF is dechlorinated, half of the DCM accumulates before its ultimate metabolism after the CF has depleted, followed by methane production (Figure 4). No other metabolites, including chloromethane, were detected accumulating in the culture.

**Figure 4.**
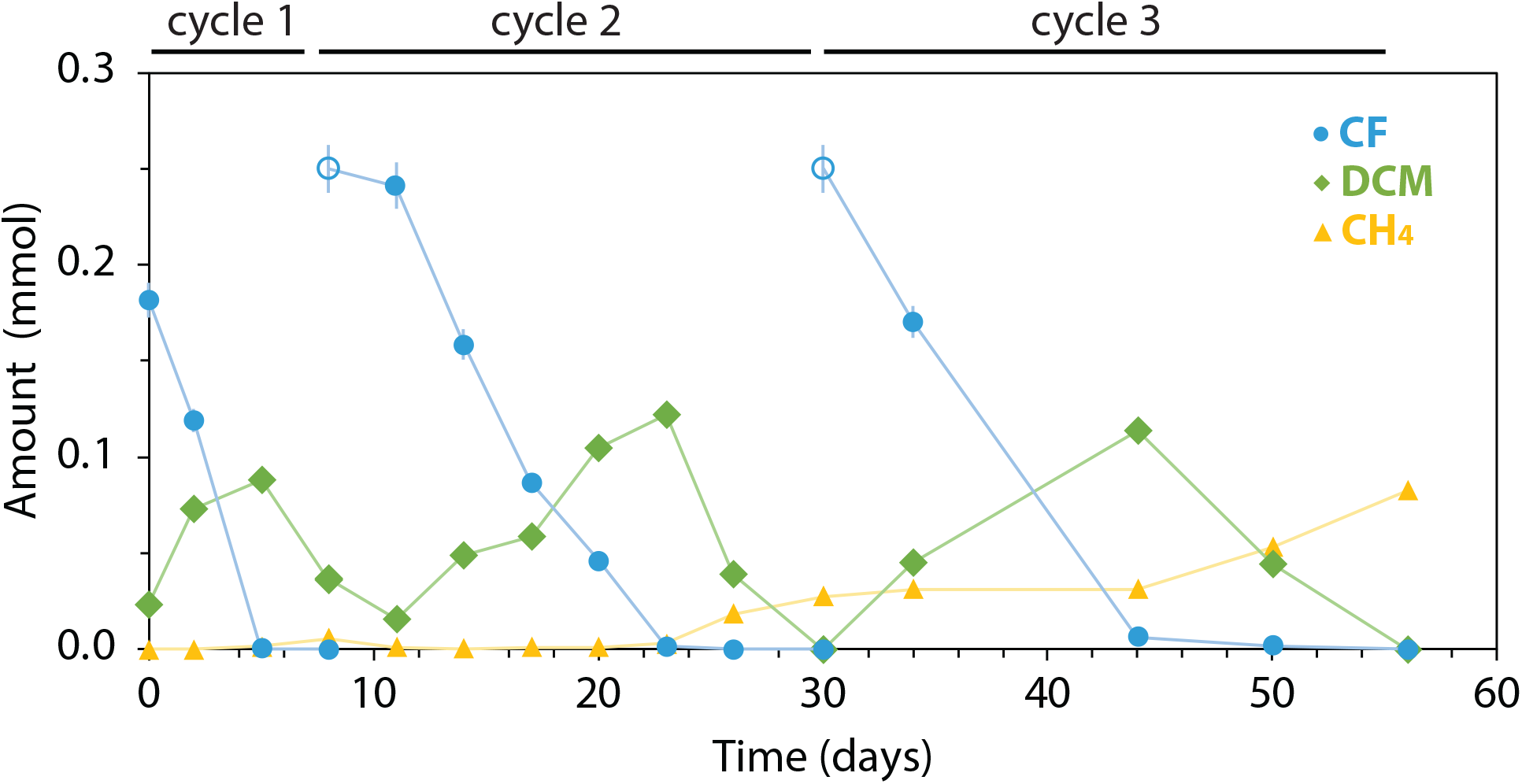
SC05-UT growth profile over three feeding cycles (Days 1327-1383, Figure 2) used for electron balance calculations (n=1 bottle, error bars indicate analytical uncertainty of each measurement). Open circles denote CF amendments.

An electron balance was performed over each of the three feeding cycles (Table 1). The first two cycles showed higher recovery of electron donor than acceptor (111% and 103%, respectively), suggesting an acceptor that was unaccounted for—likely hydrogen production during DCM mineralization. Previous work has shown that SC05 transforms CF to carbon dioxide and hydrogen (18), but the theoretical amount of hydrogen required to correct for the discrepancy in this electron balance is below the detection limit (2% of the bottle’s headspace) for our instruments.

**TABLE 1.** Electron balance in SC05-UT during CF and DCM degradation over 3 feeding cycles.

| Cycle | Duration<br>(days) | Acceptors consumed (μeeq) | | | | Donors<br>consumed<br>(μeeq) | Percent *e*-<br>recovery<br>(donor/<br>acceptor) |
| --- | --- | --- | --- | --- | --- | --- | --- |
|  |  | CF<br>dechlorination | Methanogenesis | Biomass* | SUM | DCM<br>mineralization<sup>‡</sup> |  |
| 1 | 8 | 362 | 44 | 202 | 609 | 674 | **111** |
| 2 | 19 | 482 | 211 | 308 | 1001 | 1026 | **103** |
| 3 | 22 | 340 | 411 | 259 | 1010 | 862 | **85** |
| **Total** | **56** | **1185** | **667** | **769** | **2620** | **2562** | **98** |

\*Fraction of electrons to biomass calculated from measured cell yields ( $f_s=0.3$ eeq cells/eeq donor, Table S2).
<sup>‡</sup>DCM mineralization calculated by subtracting DCM accumulated from CF dechlorinated.

The third feeding cycle resulted in an 85% recovery, likely due to increased methanogenesis—which consumes the accumulated hydrogen, rendering all major electron pools detectable (Table 1). Methanogenesis is inhibited by CF, so an immediate refeeding of CF during DCM mineralization (such as at the end of cycles one and two) will prevent this hydrogen consumption and lead to a gap in electron balance data. This gap is resolved with feeding less consistently (such as at the end of cycle 3 or using cumulative data over months (Table S4).

### *Dehalobacter* remains abundant in DCME

To identify the DCM-degrading microorganism in SC05-UT, a DCM enrichment (DCME) was created and maintained on DCM alone (Figure 2). A sample of DCME’s degradation profile is shown in Figure S1, and an overall electron balance of DCME is shown in Table S4. Throughout the enrichment process, DNA samples from both cultures (blue circles, Figure 2) were also sequenced for culture composition. After each transfer of DCME, we were surprised to see the same *Dehalobacter* ASV that dominates SC05-UT increase in relative abundance during DCM enrichment (Figure S2C, S2D). Even after enriching for 1326 days, 33% of bacterial reads belonged to this *Dehalobacter* ASV. We saw several other ASVs increase in abundance, including Peptococcaceae, Burkholderia and two Spirochaete ASVs, as well as changes in methanogenic archaeal populations after each transfer (Figure S2A, S2B).

CF is a known inhibitor of many microbes due to its similarity in shape to a methyl group, rendering it inhibitory to the growth of many homoacetogens, acetate-consuming sulfate reducers, and methyltransferase-using bacteria—including most known DCM-degrading organisms—which is a key reason for its harm to carbon cycling in subsurface ecosystems (24). The maintenance conditions of SC05-UT include frequent amendments of CF, which lead to the selection of CF-resistant organisms including *Dehalobacter*. In DCME, where this condition is lifted, those organisms are now free to proliferate in the culture, consuming organic acids in the culture or hydrogen produced from DCM mineralization. Given this freedom for faster-growing fermenters to flourish, 33% abundance is surprisingly high for the relatively slow-growing *Dehalobacter* (Figure S2D).

These results were echoed when more closely monitoring DCME’s culture composition over one feeding period (Experiment #1, Figure 2). Here, the archaeal populations remained stable while there were temporal fluctuations in abundance of multiple bacterial ASVs (Figure S3A, S3B). Over 30 days, the DCME community consumed DCM and ultimately produced acetate and methane, while no degradation or production was seen in the killed controls (Figures S3C, S4). Over the DCM transformation period, the most notable increase in fraction abundance was the *Dehalobacter* ASV, though the Peptococcaceae ASV also increased in relative abundance (Figure S3B).

Without absolute quantification, the growth (or perseverance) of any of these microbes cannot be attributed to DCM metabolism, compared to consumption of other metabolites resulting from the removal of CF’s inhibition in the system. Further experimentation was needed to establish the relationship between *Dehalobacter* and both DCM mineralization and CF dechlorination, thus additional transfers were set up (Experiment #2, Figure 2) and the community compositions were quantified.

### *Dehalobacter* dechlorinates CF and mineralizes DCM

In the Experiment #2 sub-transfers (Figure 5), not only the relative abundance, but also the absolute quantity of *Dehalobacter* was tracked during CF and DCM degradation in SC05-UT and DCME sub-transfers, respectively. The number of *Dehalobacter* 16S rRNA copies increases as CF is degraded over 41 days (Figure 5A, 5C), yielding an average of 4.92 ± 0.3 × 10^13^ cells/ mol CF, assuming three 16S rRNA copies per cell (Table 2). The SC05-UT *Dehalobacter* yield is similar to, though slightly higher than, that of other known CF-dechlorinating bacteria (Table 2). More notably, the number of *Dehalobacter* 16S copies also increases as DCM is depleted in the DCME sub-transfers (Figure 5B, 5D). The calculated yield is 1.26 ± 0.1 ×10^13^ cells/ mol DCM, which is on the same order of magnitude as other known DCM-degrading organisms, excluding *C*. Formimonas warbariya (Table 2). Additional growth on DCM may account for the slightly higher *Dehalobacter* yield when growing on CF as well, as small amounts of DCM produced from CF dechlorination may be consumed by *Dehalobacter* concurrently. No degradation or *Dehalobacter* growth was detected in the killed controls (Figure S5).

**Figure 5.**
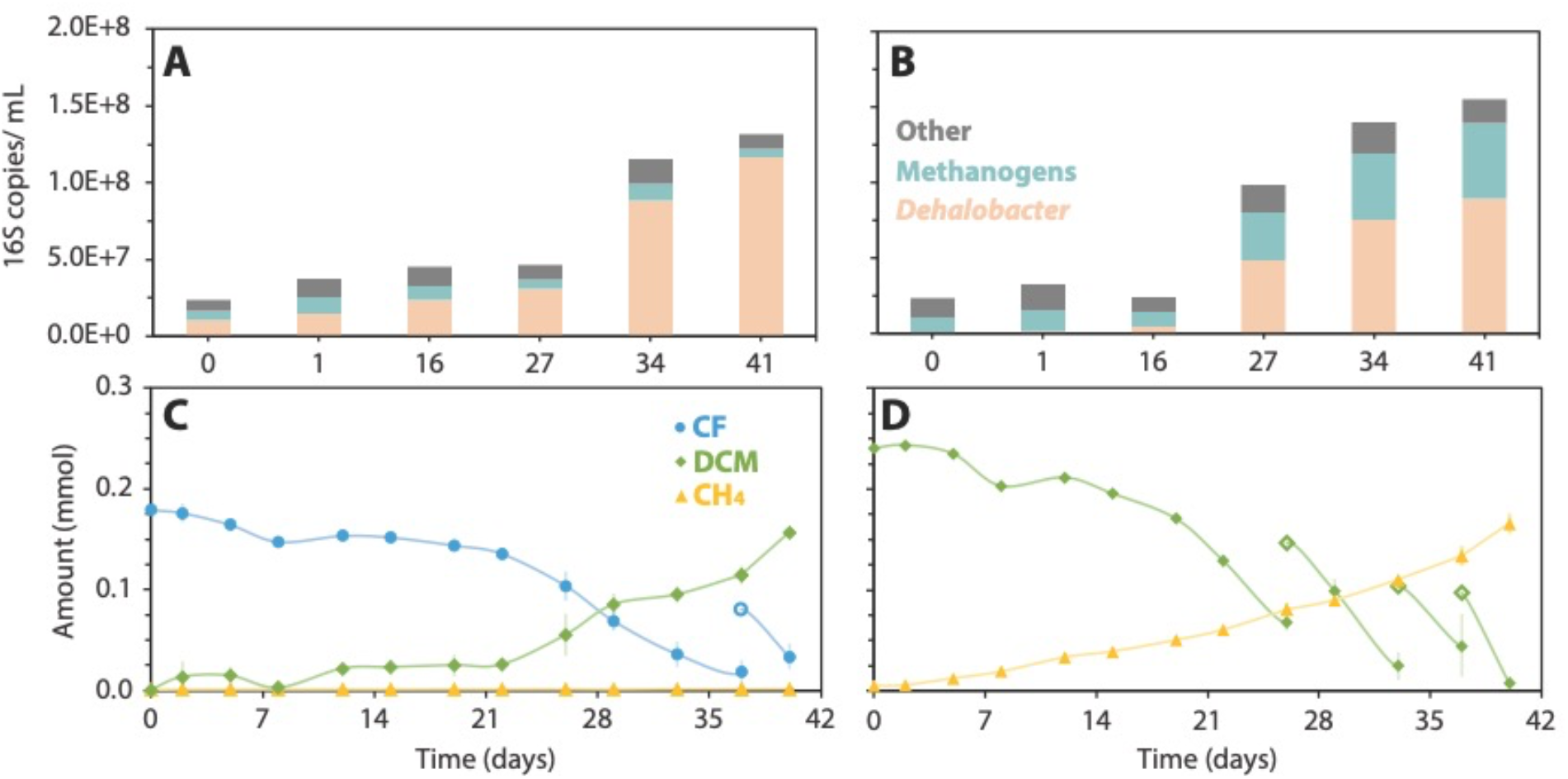
*Dehalobacter* abundance increases with CF dechlorination in SC05-UT (A, C), and DCM mineralization in DCME (B, D) over 42 days (Days 817-858, Figure 2). Panels A and B contain 16S amplicon data scaled to detected 16S rRNA copies per sample. Panels C and D show measured CF, DCM and methane in each bottle over time. Error bars represent biological variation (n=3). Open circles/ diamonds indicate refeeding of CF and DCM, respectively.

**Table 2.**
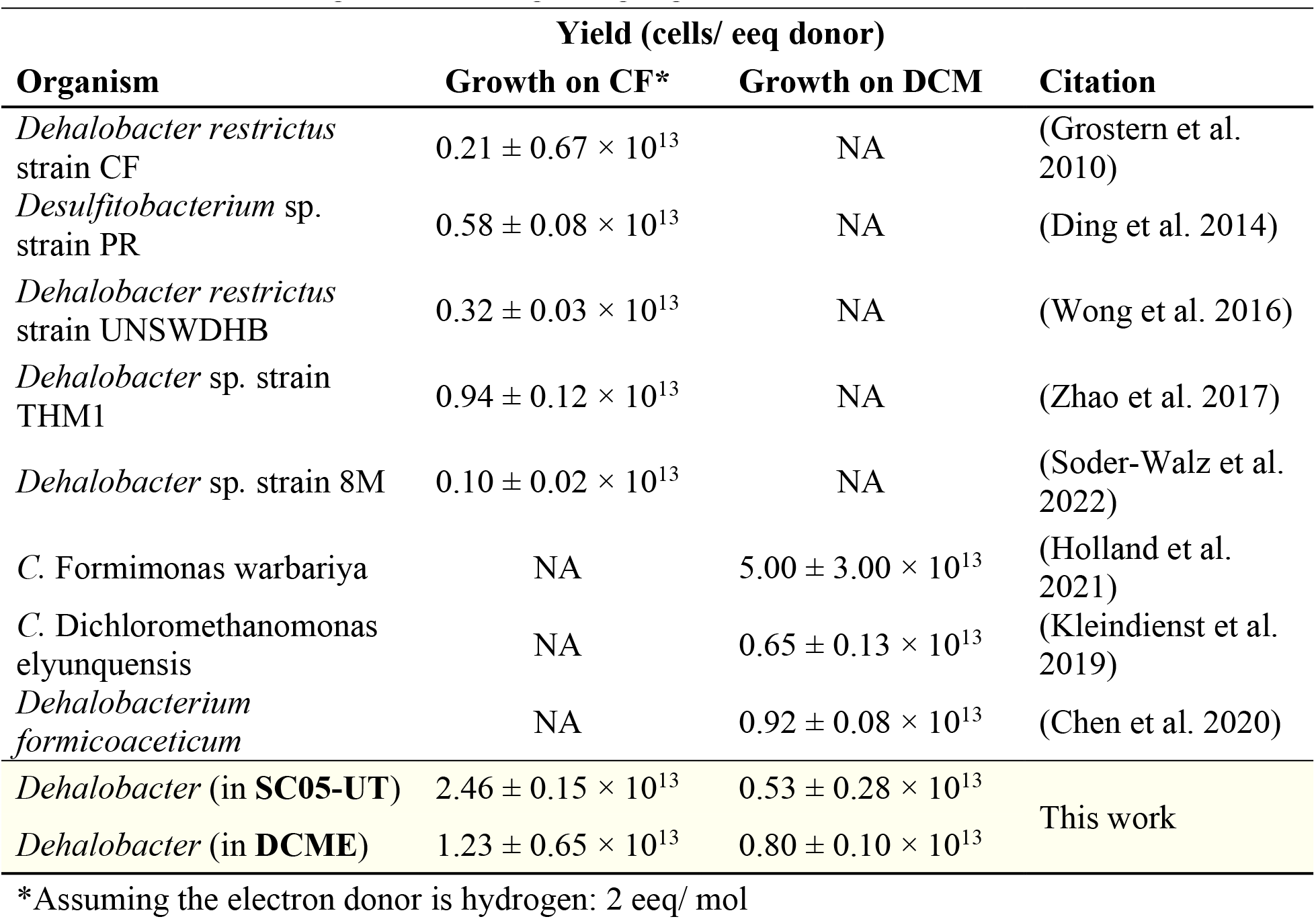
Cell yields of *Dehalobacter* when growing on CF compared to DCM alone, compared to known CF-dechlorinating and DCM degrading organisms.

In both degradation phases, the only organism to increase in abundance proportionally to the substrates’ consumption is a single *Dehalobacter* ASV (Figure 5A, 5B). DCME also shows increase in methanogen ASVs over the degradation cycle (Figure 5B), whose activity may serve as a sink for the hydrogen produced during DCM mineralization. These data suggest that either 1) a single *Dehalobacter* strain or 2) two *Dehalobacter* strains indistinguishable by 16S amplicon sequencing are performing both transformation reactions.

### A single *Dehalobacter* strain performs both transformation steps?

Cultures containing multiple strains of *Dehalobacter* that each use different sequential dechlorination products as electron acceptors have been previously reported, such as the ACT-3 culture that contains *Dehalobacter restrictus* strains CF and DCA, (25). Strain CF can dechlorinate 1,1,1-trichloroethane to 1,1-dichloroethane, which is then dechlorinated by strain DCA to chloroethane, but after enrichment of the parent culture on 1,1-dichloroethane, the culture no longer degrades 1,1,1-trichloroethane. If a similar system exists in SC05-UT, it should no longer be readily able to dechlorinate CF after enrichment on DCM. However, after 1203 days without CF, DCME maintained its ability to dechlorinate CF, which was completely depleted in 16 days (Figure 6), while killed controls showed no CF dechlorination (Figure S6). DCM, though initially accumulating, was depleted after 28 days. Hydrogen added to the bottles’ headspace used to kickstart dechlorination likely explains the lag in DCM degradation, reducing the need for DCM as an electron donor or even preventing its metabolism due to potential inhibition of hydrogen-evolving hydrogenases needed for mineralization (17).

**Figure 6.**
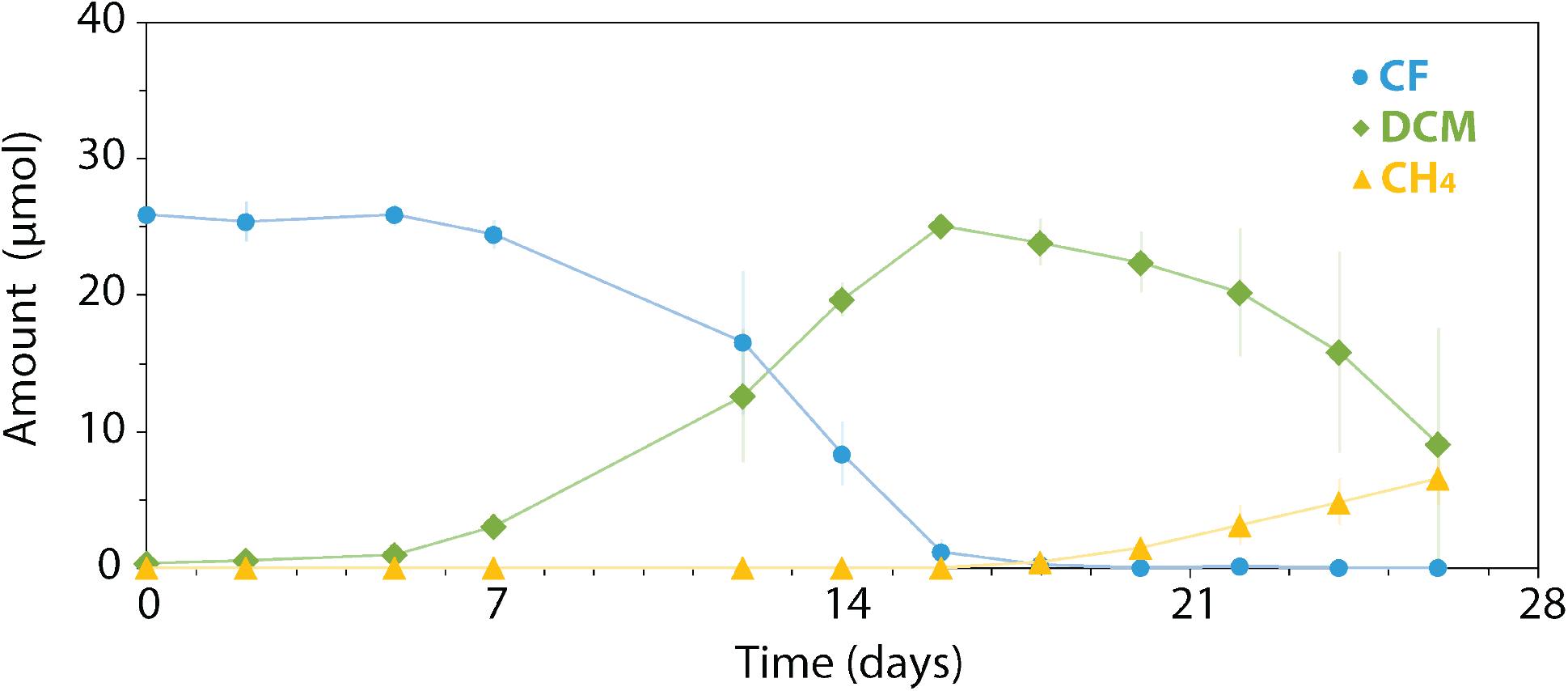
Dechlorination profile of DCME when re-amended with CF and hydrogen after 1203 days of DCM enrichment (n=2, error bars represent biological error).

The ability of DCME to quickly adapt to the addition of CF supports the proposal of a single *Dehalobacter* strain with a flexible metabolism performing both dechlorination steps in SC05-UT. There is a lag period of about seven days after feeding that may be necessary for regulation and expression of various genes related to CF metabolism or protection from CF exposure. It is worthy of note however, that SC05-UT often experiences similar lag periods after transfer to a new bottle (as seen in Figure 5C), which may be due to a disruption in the fine balance of electron transfer in the community and could explain the lag in DCME as well.

Growth of *Dehalobacter* was also tracked during this experiment (Figure S7), allowing calculation of cell yields on CF alone and DCM alone in both SC05-UT and DCME, which are compared in Table 2. The *Dehalobacter* cell yields are comparable between SC05-UT and DCME during each metabolic phase. During DCM degradation, the yields are within error of each other, while during CF dechlorination, we see a slightly higher yield in SC05-UT than DCME (Table 2). This could be due to a more streamlined concurrent degradation in SC05-UT, which is used to degrading both CF and DCM at once.

### Conclusions and Implications

Though further genomic and experimental investigation of the *Dehalobacter* strain(s?) in SC05 are necessary to confirm whether a single strain or multiple closely related strains are responsible for tandem CF and DCM dechlorination, this study shows the importance of *Dehalobacter* in each transformation step. We also show the possibility for intraspecies or intracellular electron transfer to circumvent the need for exogenous electron donor in the culture. Typical bioaugmentation cultures for dechlorination require the co-injection of electron donors to provide a source of reducing equivalents for reductive dechlorination to occur. This costly process is one of the most financially prohibitive steps of the bioaugmentation process, thus favoring a culture that circumvents this requirement, such as SC05. This culture is already in use commercially, marketed as KB-1® Plus CF, and with a clearer understanding of its electron donor requirements, or lack thereof, it could allow for more efficient and less expensive groundwater remediation worldwide.

## ASSOCIATED CONTENT

### Supporting Information

Growth profile of DCME; SC05-UT and DCME community compositions during enrichment; DCM degradation community composition and metabolite profile; Negative controls for Experiment #1; Negative controls for Experiment #2, Negative controls for Experiment #3; *Dehalobacter* yield during CF degradation by DCME (PDF)

Staggered primer sequences; Electrons to biomass calculations and assumptions; Yield calculations; Cumulative electron balance in SC05-UT and DCME (XLSX)

## AUTHOR INFORMATION

## Author Contributions

E.A.E., R.M., and O.B. conceived of the project. The original SC05 culture was established by J.W. and S.D. SC05-UT and DCME were established and maintained by O.B. O.B. established all experimental bottles, and performed all GC and IC measurements, DNA extractions, qPCR reactions, and amplicon sequencing data analysis. Electron balances and yield calculations were performed by O.B. and E.A.E. The manuscript was written by O.B., R.M. and E.A.E. All authors contributed to manuscript revision and have approved the submitted version.

## ACKNOWLEDGMENTS

This work was funded by the Natural Science and Engineering Research Council (NSERC) though a Discovery Grant to E.A.E. and a Doctoral Scholarship to O.B., as well as by a Genome Canada Bioinformatics and Computational Biology (BCB project 285MPR) subgrant to R.M. and E.E. We would also like to thank SiREM for providing us with the culture studied in this work, as well as Elizaveta Saitova helping with data collection, and Elizabeth Phillips for her valuable insights into caring for the culture.

