## Supplementary Information for "A multifunctional *Dehalobacter*? Tandem chloroform and dichloromethane degradation in a mixed microbial culture"

**Supplementary Tables (in accompanying Excel file)**

**Table S1.** Staggered primers used for amplicon sequencing.

**Table S2.** Calculation of fraction of electrons to biomass (fs) based on thermodynamic calculations and experimental values.

**Table S3.** *Dehalobacter* yield calculations on CF and DCM.

**Table S4.** Cumulative electron balance in SC05-UT and DCME.

**Supplementary Figures**

**Figure S1.** Cumulative DCM dechlorinated by the DCM enrichment and the overall growth profile over 830 days.

**Figure S2.** Community composition of SC05 parent culture and DCM enrichment throughout enrichment.

**Figure S3.** Community composition and metabolite profile of a DCM enrichment subtransfer over one feeding cycle

**Figure S4.** DCM, acetate, and methane in negative controls for Experiment #1.

**Figure S5.** CF and DCM measured in negative controls and copies of *Dehalobacter* 16S rRNA in for Experiment #2.

**Figure S6.** CF, DCM, and methane in negative controls for Experiment #3.

**Figure S7.** *Dehalobacter* yield in DCME compared to dechlorination profile when re-amended with CF.

**References for SI**

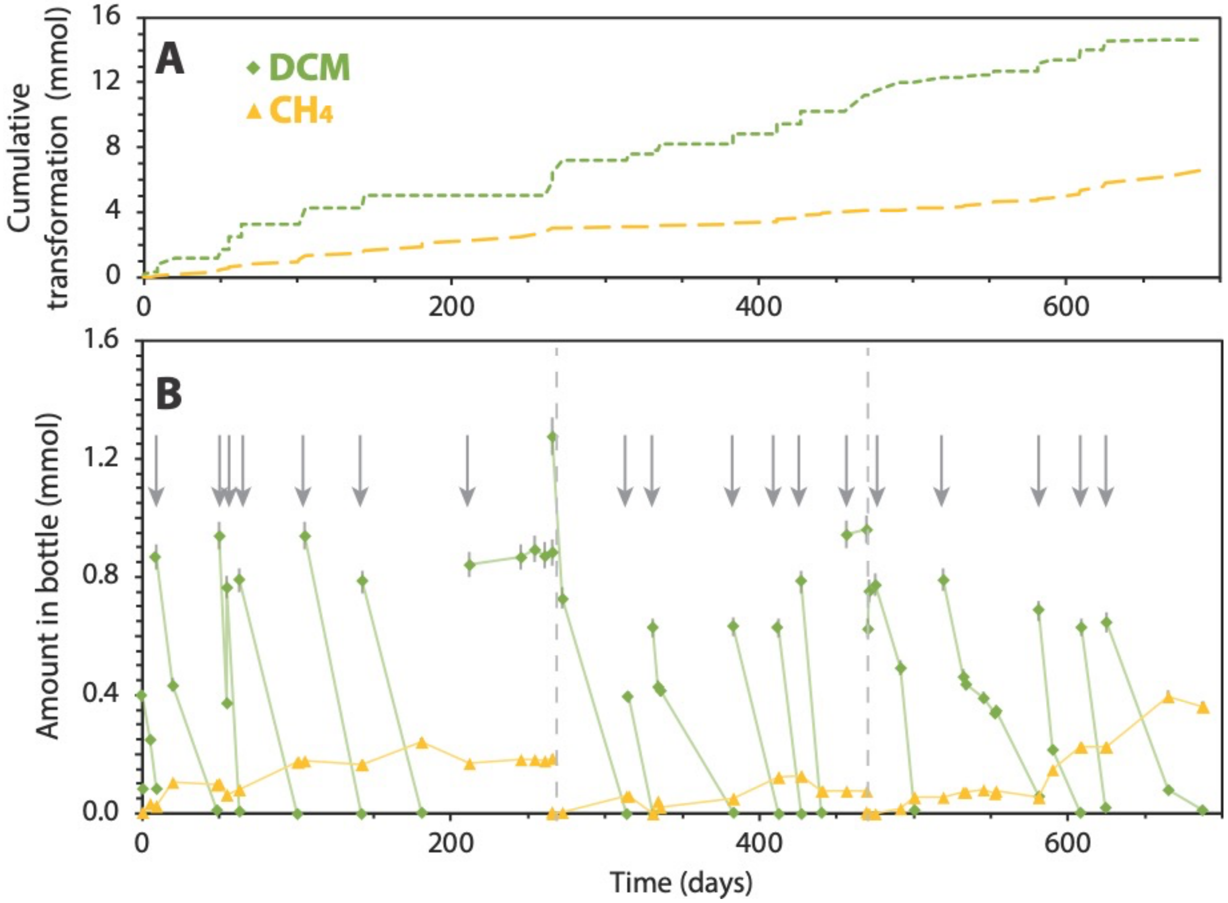


**Figure S1. A)** Cumulative DCM dechlorinated and methane produced by DCME (Transfer 3, Days 482–1170 in Figure 0.5) and **B)** the overall growth profile over 688 days. Arrows indicate refeeding of DCM, and grey dashed lines denote headspace purging of methane from bottle (n=1, error bars indicate analytical uncertainty of each measurement).

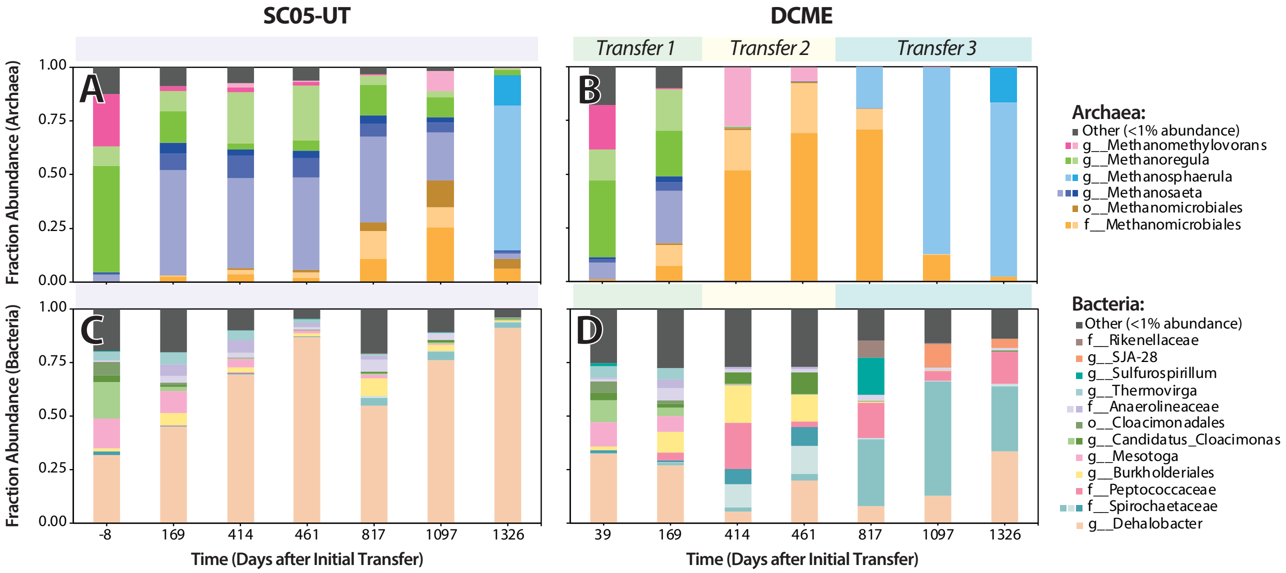


**Figure S2.** Community composition of **A, C)** SC05-UT and **B, D)** DCME over 1326 days (sampling points shown in Figure 0.5). Percent abundances of archaeal reads are shown in panels A and B, while bacterial reads are shown in panels C and D. Each colour represents one ASV.

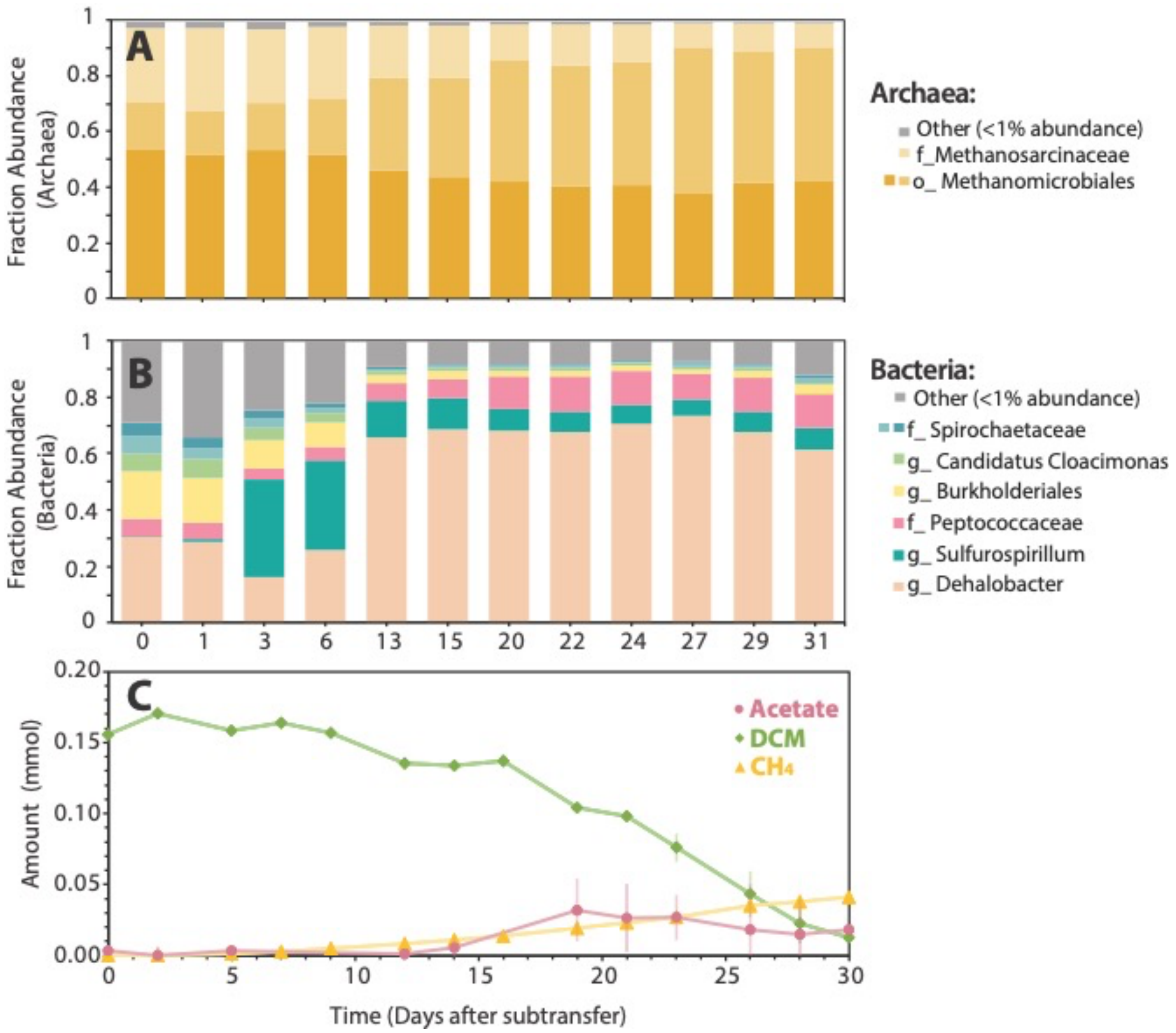


**Figure S3. A)** Archaeal and **B**) bacterial community compositions, and **C)** metabolite profile of DCME sub-transfers over one feeding cycle (Experiment #1, Days 433-463, Figure 0.5). Error bars represent biological error (n=3). Each colour represents one ASV.

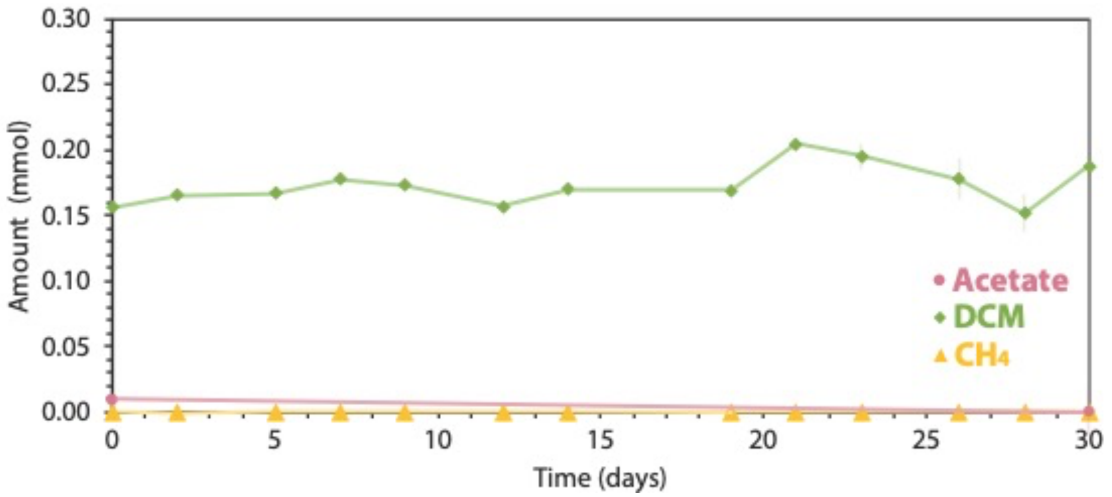


**Figure S4.** DCM, acetate, and methane in negative controls (killed culture) for Experiment #1 (n=3). Error bars denote replicate error.

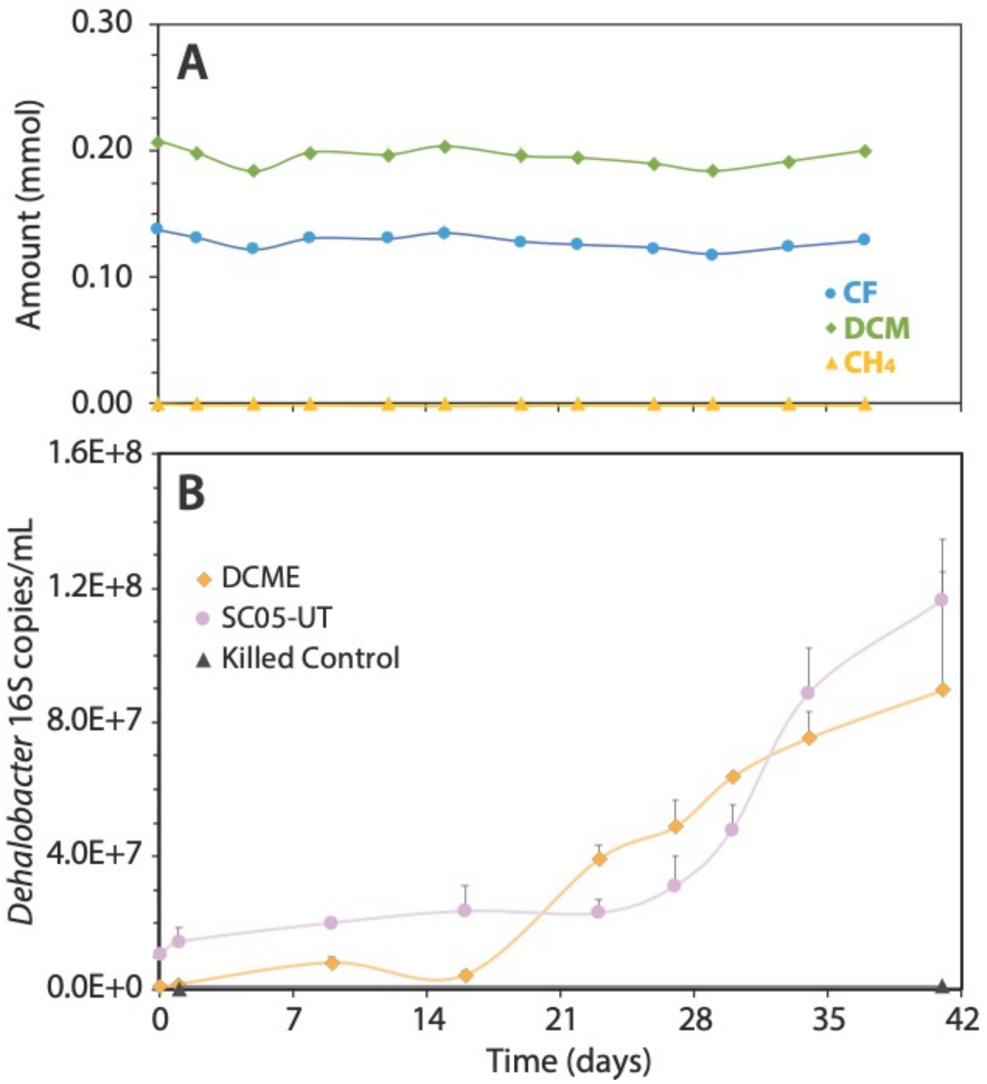


**Figure S5. A)** CF and DCM measured in negative controls (killed culture) during Experiment #2 (n=2) and **B)** copies of *Dehalobacter* 16S rRNA in DCME, SC05-UT and killed control. Error bars denote replicate error.

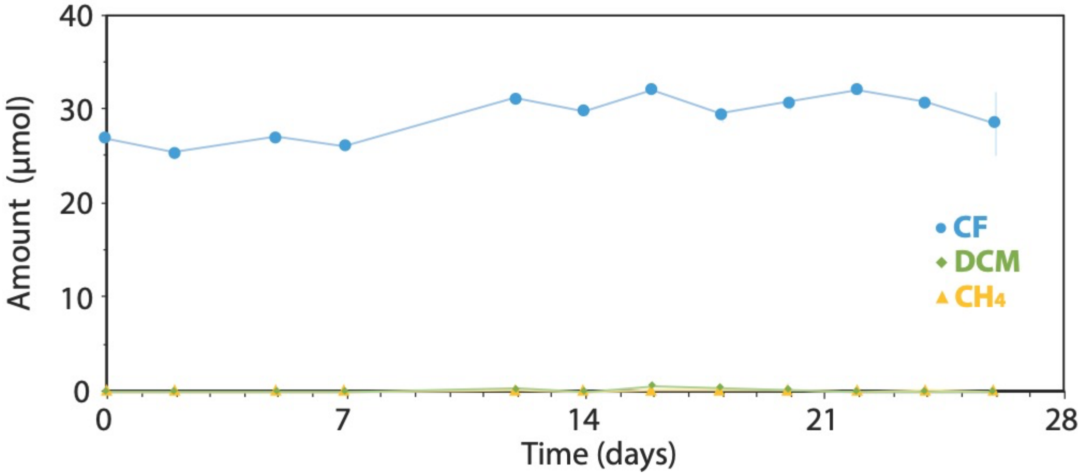


**Figure S6.** CF, DCM, and methane in negative controls (killed culture) for Experiment #3 (n=2). Error bars denote replicate error.

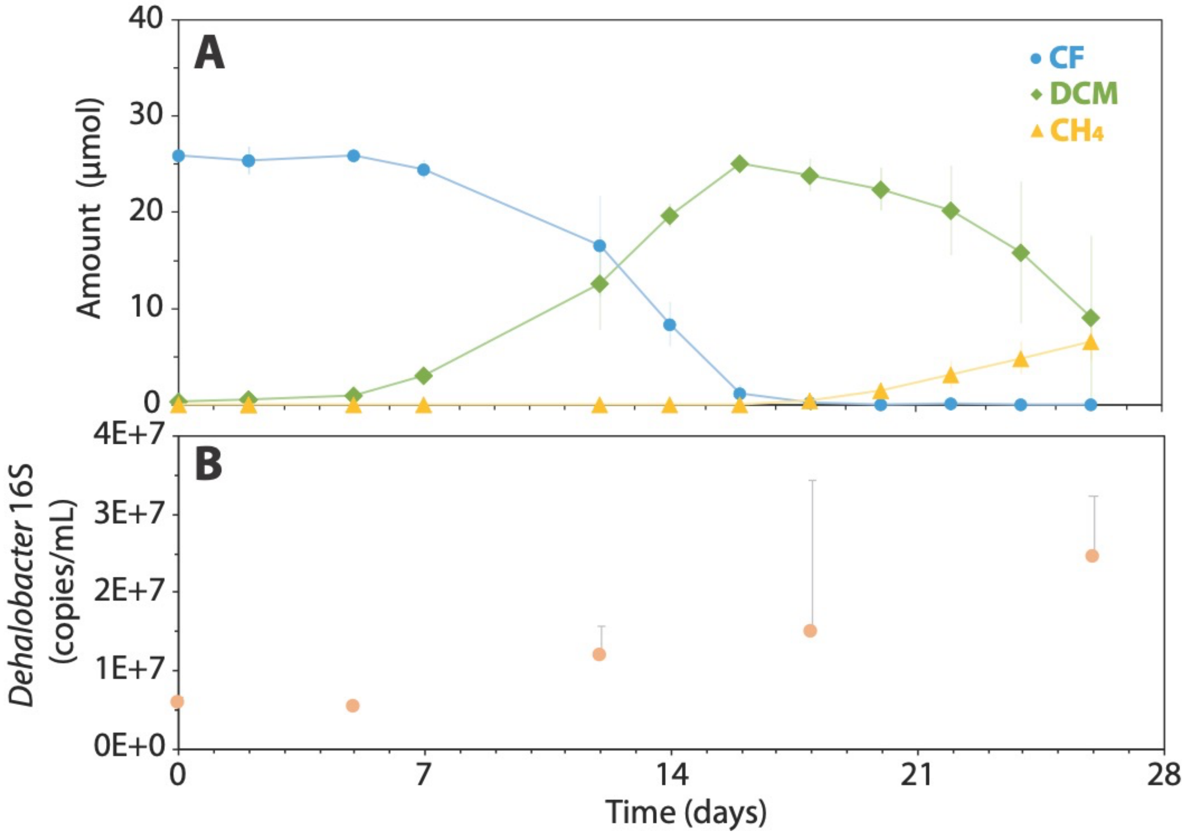


**Figure S7. A)** *Dehalobacter* yield in DCME compared to **B)** dechlorination profile when re-amended with CF during Experiment #3 (n=2, error bars represent biological error).
